# The tectum transversum(TTR) maintains patency of the developing coronal suture

**DOI:** 10.1101/2025.03.30.646197

**Authors:** Meenakshi Umar, Garrett Bartoletti, Dimitri Sokolowskei, Nathan Janser, Robert Tower, Fenglei He

## Abstract

Craniosynostosis is a congenital defect characterized by the premature fusion of calvarial bones, often attributed to the loss of fibrous sutures or deregulated bone formation. Recent studies have reported abnormal cartilage formation in multiple synostosis models, suggesting a potential role for cartilage in suture formation and maintenance. The tectum transversum (TTR) is a transient cartilage located between the coronal suture and dura, adjacent to the frontal and parietal bones. Abnormal TTR formation has been observed in several models; however, its role in coronal suture development remains unclear. In this study, we investigated the developmental process of TTR in a mouse model and characterized its formation in relation to adjacent tissues, including the calvarial bones and the coronal suture. Through genetic ablation of TTR, we demonstrated its essential role in maintaining coronal suture patency. Furthermore, spatial transcriptomics data suggest that TTR may function as a barrier to BMP signaling activation in the coronal suture, a process potentially influenced by the dura. These findings provide new insights into the mechanisms regulating coronal suture development and the etiology of coronal synostosis.

## Introduction

Craniosynostosis is a prevalent birth defect characterized by the premature closure of calvarial sutures, which normally separate and connect individual calvarial bones^1,2^. Synostosis at the coronal suture, which separates frontal and parietal bones, counts for approximately 25% of cases^3,4^. While previous studies utilizing clinical genetics and mouse models have identified various genetic factors associated with both normal coronal suture development and synostosis^3,5-7^, there still exists a significant gap of knowledge in understanding the fundamental mechanisms underlying suture development and pathogenesis, particularly at the cellular level.

Although most bones of the mammalian skull form through intramembranous ossification—distinct from endochondral ossification, which involves cartilage-a cartilaginous skull is initially established before intramembranous bones develop. This cartilaginous structure, known as the cranial endoskeleton, includes the cartilages that make up the chondrocranium and pharyngeal skeleton^8^. These cartilages typically follow one of two paths: they either undergo endochondral ossification or are resorbed^8,9^. Despite its close association with the dermal skeleton, the cranial endoskeleton’s role in craniofacial development and malformation has received relatively little attention^8,10^. However, recent studies have reported abnormal cartilage formation in multiple calvarial sutures preceding their premature fusion^11-17^. These findings highlight the increasing significance of these cartilages in craniofacial development and suggest their potential involvement in the etiology of craniosynostosis, which remains to be fully elucidated.

In previous study, we have reported that augmented PDGFRα activity leads to premature closure of coronal suture, accompanied by abnormal expansion and ossification of a cartilage anlage known as the tectum transversum (TTR), which is located between the coronal suture and underlying dura during calvarial development^8,15^. However, it remains unclear how the TTR is developed and regulated, and whether it is important for coronal suture formation.

In the present study, we thoroughly examined the formation of the TTR, tracing its development from initial emergence to eventual disappearance, while elucidating its lineage and differentiation processes. Using a genetic model, we found that ablation of the TTR leads to premature fusion of the coronal sutures. Spatial transcriptomics data reveal that the TTR preserves coronal suture progenitor cells patency by preventing their premature differentiation into osteoblasts, a process mediated through the activation of BMP signaling and cell-cell communication. These findings provide insights into the mechanisms underlying TTR development and its impact on coronal suture formation, while also shedding light on the fundamental mechanisms by which cartilage regulates calvarial suture development.

## Results

### Development of TTR during mouse embryogenesis

To investigate the developmental progression of the TTR, we conducted wholemount staining and histological analyses of mouse embryos at various stages. At embryonic day (E) 13.5, alcian blue staining revealed the emergence of TTR as a cartilage anlage above the eye (Fig. 1A). By E14.5, the developing TTR began to establish connections with other calvarial cartilages (Fig. 1B). From E16.5 to E18.5, the TTR remained positioned between the frontal bone (fb) and parietal bone (pb) but gradually shrinks in size (Fig. 1C, D). At postnatal day (P) 5, the TTR was no longer detectable (Fig. 1E, E’). Alkaline phosphatase/alcian blue (AP/AB) staining on transverse sections of mouse embryos at these stages corroborated these findings, confirming the location of the TTR beneath the coronal suture (cs) and adjacent to the fb and pb (Fig. 2). The TTR exhibited a characteristic waterdrop-like shape, with a narrow tip on the dorsal end (Fig. 2A, C, E, and G) and a wider base on the ventral end (Fig. 2B, D, F, and H). Furthermore, it was observed that the TTR experienced quick growth from E13.5 to E16.5 (Fig. 2A-F), followed by a subsequent degeneration, as evidenced by its smaller size at E18.5(Fig. 2G-I). At P5, the dorsal portion of the TTR is not detectable, and the ventral portion is replaced by ossified tissue (Fig. 2G-J).

**Figure 1.**
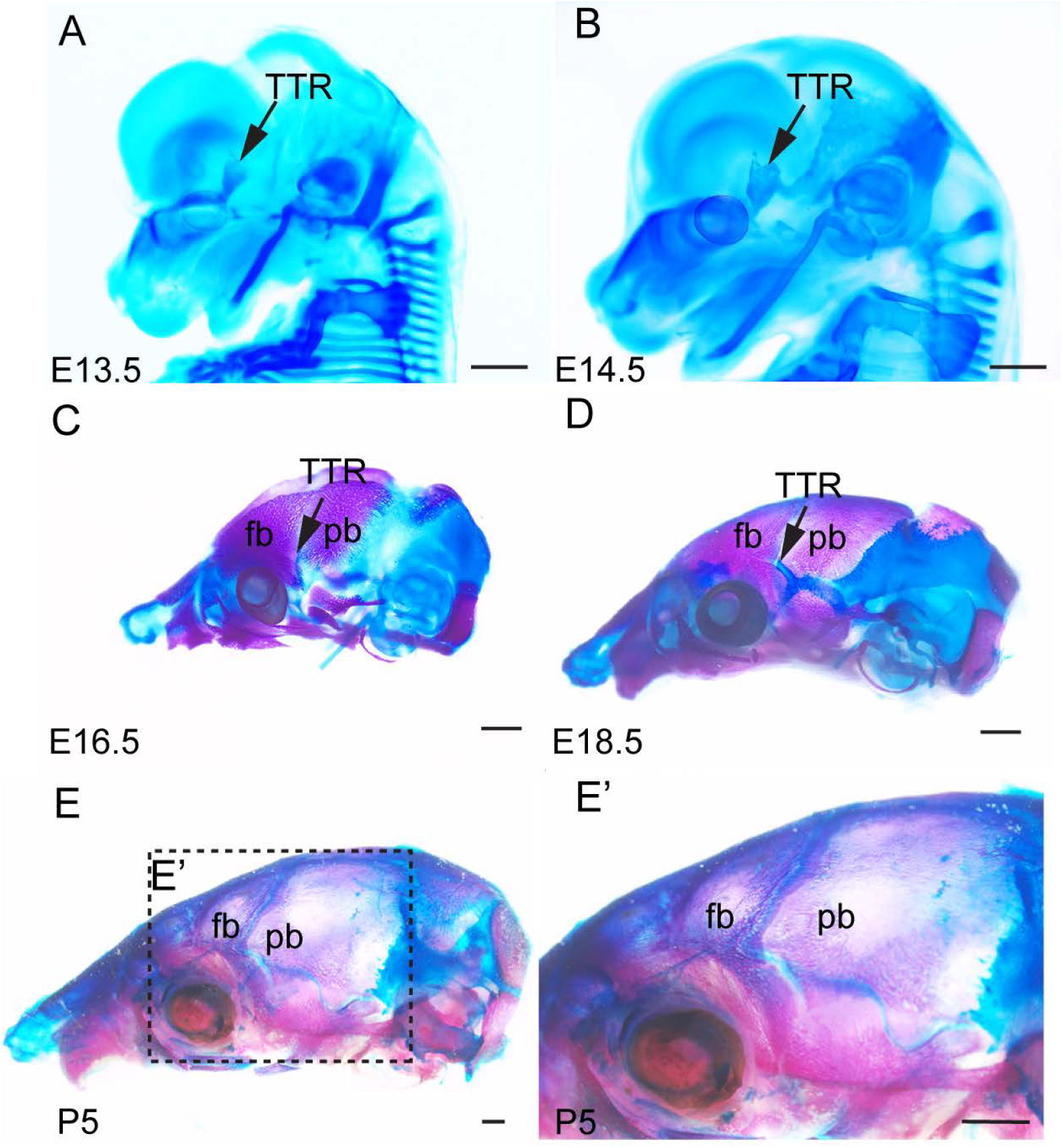
TTR morphogenesis during mouse embryo development. (A, B) Wholemount alcian blue staining of mouse embryo at E13.5(A) and E14.5 (B). (C, D, E) Skeletal preparation of mouse skull at E16.5 (C), E18.5 (D) and postnatal day (P) 5 (E). Arrows point to TTR. Scale bar, 1mm. fb, frontal bone; pb, parietal bone; TTR, tectum transversum.

**Figure 2.**
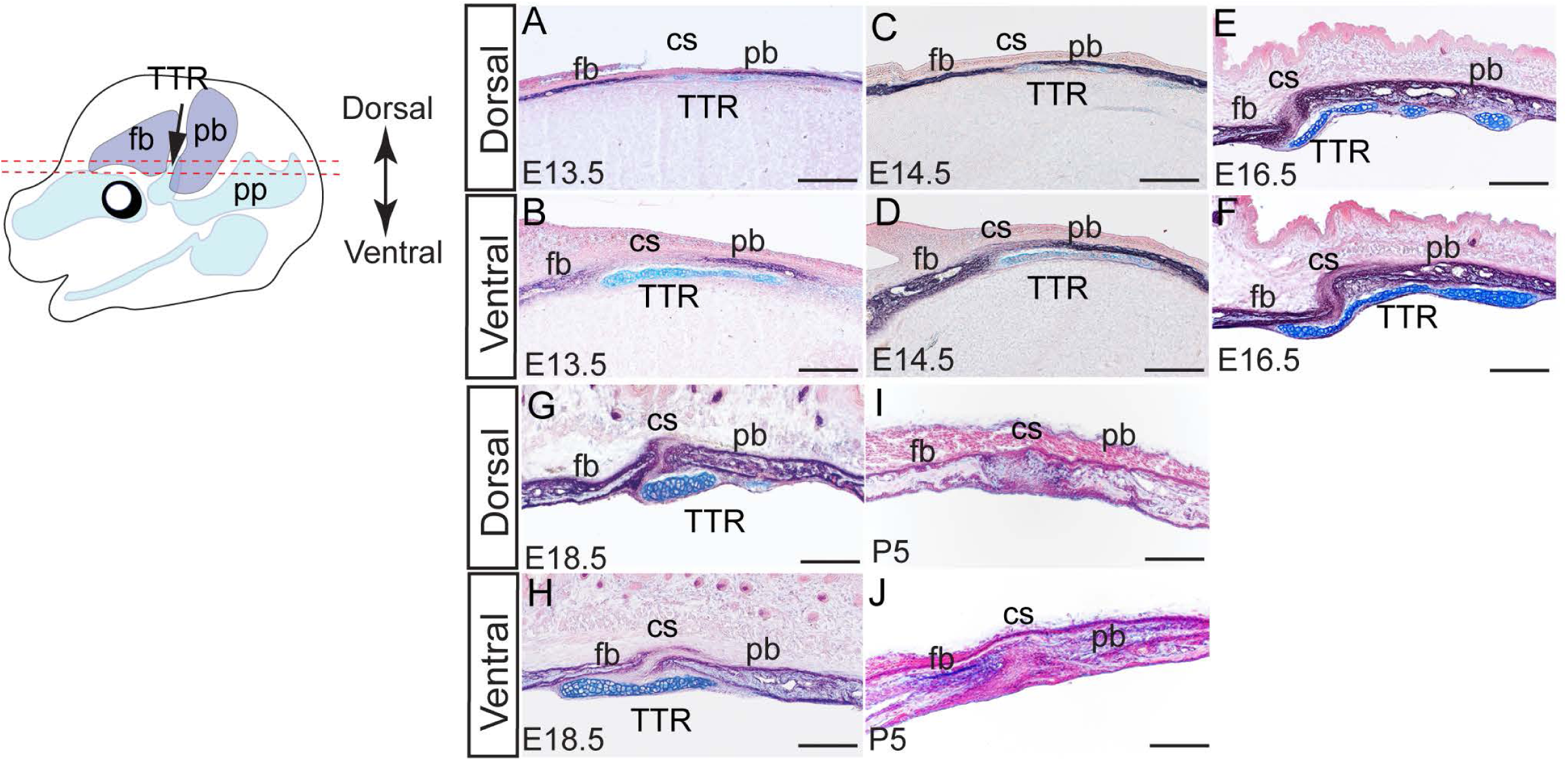
Histological analysis of the developing TTR. AP-AB staining on transverse sections of mouse heads at E13.5(A, B), E14.5(C, D), E16.5(E, F), E18.5(G, H) and P5(I, J). Sections at each stage are presented at both dorsal and ventral level. Scale bar, 100um. cs, coronal suture; fb, frontal bone; pb, parietal bone; TTR, tectum transversum.

### Lineage tracing of developing TTR

During craniofacial development, most tissues are derived from neural crest cells, with others from the mesoderm^18-20^. Of these, chondrocytes can be from either lineage. To trace the origin of TTR, we generated *Wnt1Cre2;R26R* and *Mesp1Cre;R26R* mice^21,22^. In these models, neural crest-derived tissues and mesoderm derivatives are labeled by LacZ expression, respectively. LacZ staining results showed that at E14.5, the chondrocytes of the TTR are labeled predominantly by *Mesp1Cre*(Fig. 3A, A’, B, B’), with only few cells labeled by *Wnt1Cre2* at the ventral end(Fig. 3C, C’, D, D’). This pattern remains constant to E18.5, with most TTR cells labeled by *Mesp1Cre* and only a few cells labeled by *Wnt1Cre2*(Fig. 3E-H). These results show that the TTR originated primarily from the mesoderm, with only a small contribution from neural crest at the ventral portion. This pattern is consistent with that of cs, which also originated primarily from the mesoderm^23^.

**Figure 3.**
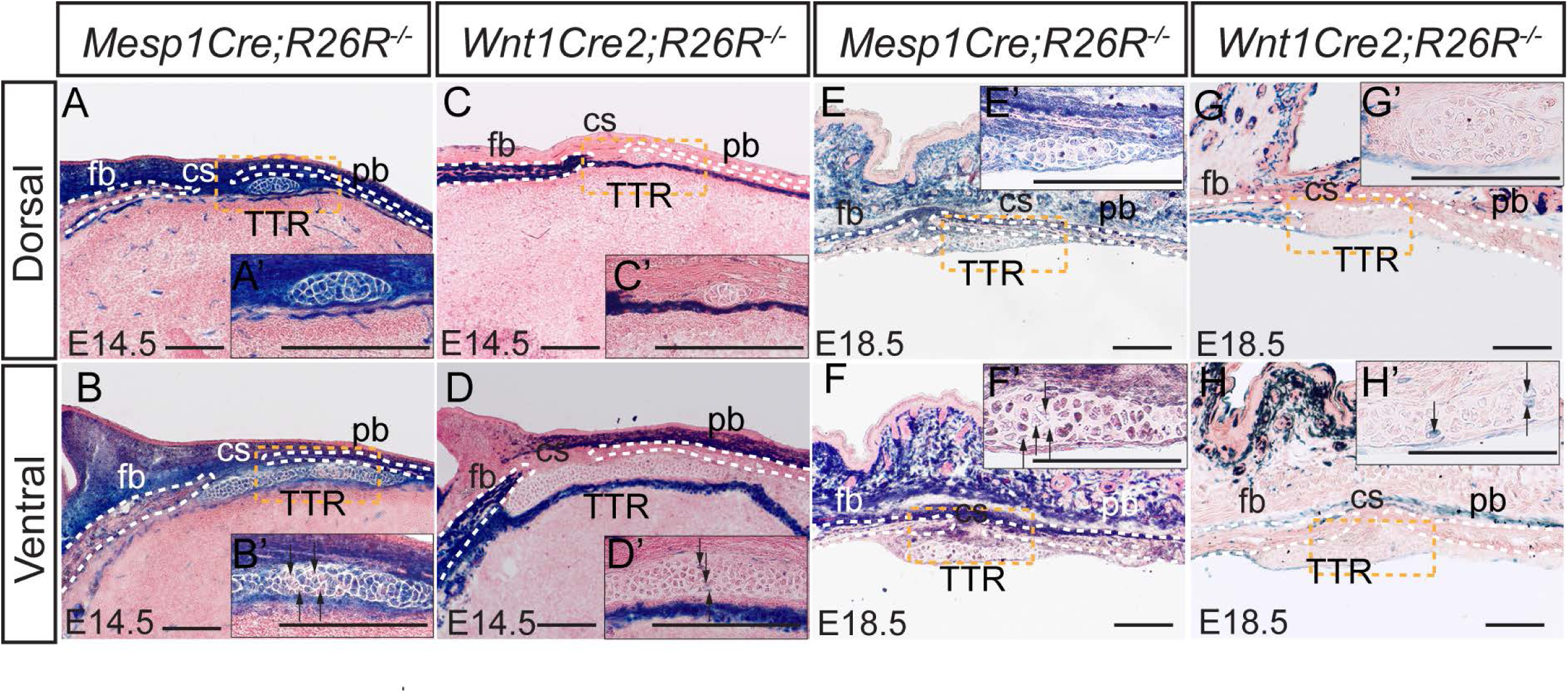
Lineage tracing of TTR. X-gal staining on transverse sections of *Mesp1Cre;R26R^-/-^* (A, B, E and F) and *Wnt1Cre2;R26R^-/-^* (C, D, G and H) at E14.5(A-D) and E18.5(E-H). Cells derived from each lineage are stained blue. The slides were then counterstained with nuclear fast red. Insets show amplified image of area outlined by dashed lines. Scale bar, 100um. cs, coronal suture; fb, frontal bone; pb, parietal bone; TTR, tectum transversum.

Our data suggest that TTR is a transient cartilaginous tissue during development. It is thus interesting to investigate the fate of TTR chondrocytes. Chondrocytes differentiate into hypertrophic chondrocytes, then adopt one of two pathways: to undergo apoptosis or to trans-differentiate into osteoblasts^24,25^. Our previous study showed that TTR cells starts to express *Col10a1* at E15.5^15^. Our immunofluorescence staining data show that ColX and Runx2 are expressed in TTR chondrocytes at E16.5 through E18.5(Fig. 4A-D). Alcian blue staining show these chondrocytes are hypertrophic at E18.5(Fig. 4E). All these data indicate TTR chondrocytes differentiation into hypertrophic chondrocytes during embryonic development. To test whether these hypertrophic chondrocytes undergo apoptosis, we examined presence of cleaved caspase 3, a hallmark of apoptosis. No notable cleaved caspase 3 expression was observed at E15.5 and E18.5 suggesting that these cells take alternative pathway (supplemental Fig.1). To test whether these hypertrophic cells transdifferentiate into osteoblasts, we have examined expression of genetic marker for osteoblast: Col1a1 and Osx. Positive results from immunostaining (Osx) and in situ hybridization (*Col1a1*) demonstrated that both are present in the TTR(Fig. 4F, G). In addition to these osteogenic markers, the TTR was also found to express multiple genes implicated in transdifferentiation of hypertrophic chondrocytes into osteoblasts, including *Mmp2*, *Mm13* and *Mmp14*(Fig. 4H, H’, I, I’, J and J’)^25,26^. Of these, Mmp13 expression is found upregulated in hypertrophic chondrocytes turning into osteoblasts^25^, Mmp14 is essential to cleave PTH1R in chondrocytes derived osteoblasts^26^, and Mmp2 has been reported to be important for Mmp3 activation^27^. Together, these data suggest that TTR chondrocytes may preferentially undergo transdifferentiation during embryonic development.

**Figure 4.**
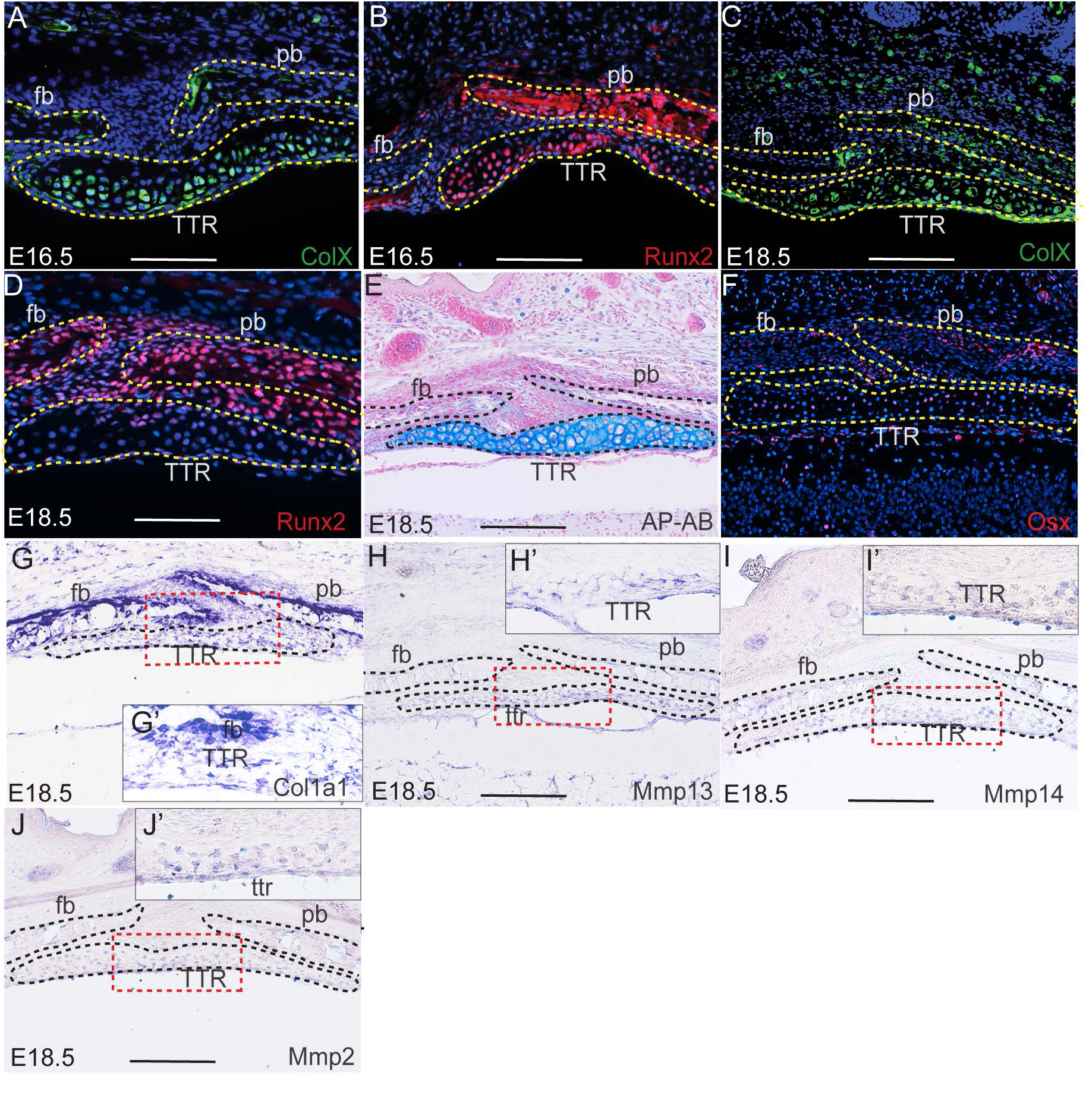
Differentiation of TTR chondrocytes. (A-D) Immunostaining on transverse sections of wildtype mouse embryos at E16.5 (A, B) and E18.5(C, D) with anti-ColX and anti-Runx2 antibody, respectively. The slides were then counterstained with DAPI(blue). (E) AP-AB staining on transverse section of E18.5 embryonic heads across TTR. (F) Immunostaining on transverse section of wild type embryonic head at E18.5 with anti-Osx antibody. (G-J) In situ hybridization with antisense probe of *Col10a1* (G), *Mmp13* (H), *Mmp14* (I) and *Mmp2* (J). Insets show amplified image of area outlined by dashed lines. Scale bar, 100um. cs, coronal suture; fb, frontal bone; pb, parietal bone; TTR, tectum transversum.

### TTR is essential to maintain cs during mouse development

The TTR is formed between the coronal suture and dura along the dorsal-ventral axis, and between the frontal and parietal bones along the anterior-posterior axis. Such a location suggests a potential role for the TTR in regulating the tissues responsible for suture formation, maintenance and/or closure. Our previous study reported abnormal enlargement and ossification of the TTR and coronal synostosis in *Pdgfra^+/K^;Meox2Cre* mice^15^. To illustrate the role of the TTR in cs formation and maintenance, we have generated *Col2a1Cre;R26R^DTA^* mice, in which Col2a1+ derived cells, including the TTR chondrocytes, are ablated by expression of diphtheria toxin A (DTA). Our data show that *Col2a1Cre;R26R^DTA^* mice do not survive beyond P0, consistent with previous report^28^. Successful ablation of cartilages in *Col2a1Cre;R26R^DTA^*at E14.5 was shown by wholemount AB staining(Fig. 5A, B), and is confirmed by AP/AB staining on transverse sections(Fig. 5A’, B’, A’’ and B’’). At E16.5, the frontal and parietal bones of *Col2a1Cre;R26R^DTA^* became thicker and started to fuse at the ventral level next to the dura(Fig. 5D, D’, and D’’), while the control counterparts remain separated(Fig. 5C, C’ and C’’). At E18.5, the frontal and parietal bones remain separated by the cs in control embryos (Fig. 5E, E’ and E’’). In *Col2a1Cre;R26R^DTA^*, these bones mostly fused at the ventral level(Fig5. F, F’ and F’’). These data indicate that the TTR is required to maintain a patent cs. The fusion between the frontal and parietal bones starting from the basal level near the dura, suggesting a potential impact of the dura on coronal suture formation in the absence of the TTR.

**Figure 5.**
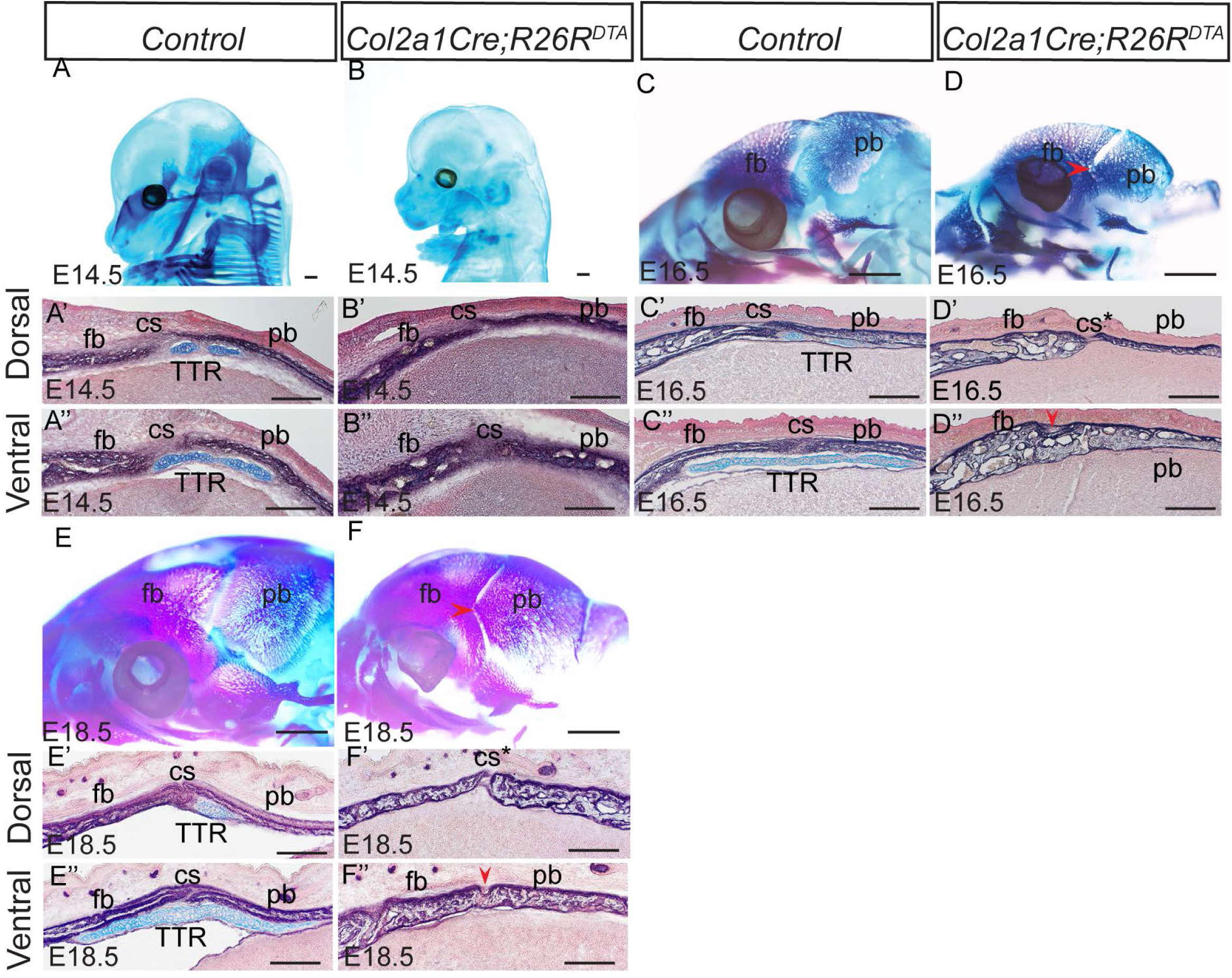
TTR is required to maintain CS in mouse embryos. (A, B) Wholemount alcian blue staining of wildtype littermate control (A) and *Col2a1Cre;R26R^DTA^* (B) at E14.5. (A’, A’’, B’, B’’) AP-AB staining on transverse section of A and B across TTR at dorsal and ventral level. Slides were counterstained with nuclear fast red. (C, D, E and F) Skeletal preparation of wildtype littermate control (C, E) and *Col2a1Cre;R26R^DTA^* (D, F) at E16.5. (C’, C’’, D’, D’’, E’, E’’, F’ and F’’) AP-AB staining on transverse section of C-F across TTR. Red arrowheads in D’’ and F’’ point to fusion sites in *Col2a1Cre;R26R^DTA^*. Scale bar in A-F, 1mm. Scale bars in A’-F’ and A’’-F’’, 100um. cs, coronal suture; fb, frontal bone; pb, parietal bone; TTR, tectum transversum.

### Spatial transcriptomics analysis on developing CS and adjacent tissues

To elucidate the molecular mechanism of how the TTR regulates the cs and adjacent calvarial bones, we performed 10X Visium spatial transcriptomics on *Col2a1Cre;R26R^DTA^* and *Col2a1Cre* control littermates. Transverse sections across the TTR of E14.5 embryos were used owing to the fact that the cs remains patent and comparable in both control and mutant embryos, and immediately precedes cs fusion in mutant mice (Fig. 5). At this stage, the TTR, fb, pb, dura and cs can be identified by their characteristic morphology (Fig. 6A). The distribution of these tissues was further confirmed by genetic markers, for example, Comp for cs, Bglap for osteo, Zeb1 for dura and Mkx for TTR (Fig. 6B). Pathway analysis of differentially expressed genes revealed that in the wt cs, genes linked to ECM organization and connective tissue development are enriched, followed by cartilage development and bone morphogenesis. In contrast, mutant cs showed enrichment in genes involved in Rho signal transduction, negative regulation of development process and positive regulation of cell differentiation (Fig 6C). Since our data show ablation of the TTR leads to malformation of adjacent tissues, such as bony fusion and impaired cs patency, we next carried out cell communication analysis. Our data show that in wt, there is active and reciprocal overall communication between the TTR and the other three tissue compartments. In the mutant, however, with ablation of TTR, communication derived from the dura signaling to the cs (blue line) is notably enhanced(Fig 6D). Specifically, we identified increased BMP signaling from the dura to the cs present within the mutant tissue, while no BMP signaling was predicted to occur in control mice (Fig 6E). Further evaluation of the BMP pathway identified upregulation of signaling mediated by dura-derived Bmp5, 6, 7 activating their receptors Bmpr1a, Acvr1 and Acvr2a present within the cs(Fig 6F). Consistent with this increased ligand/receptor signaling, BMP signaling (defined as the normalized expression of a list of gene all previously annotated to be involved in BMP signaling cascade activation) substantially increased in mutant osteo and cs tissue (Fig 6G). Conversely, we noted genes encoding BMP signaling inhibitors, including Fstl1 and Inhba, were preferentially expressed in wt TTR and cs, respectively (Fig 6H), providing a likely mechanism through which BMP signaling is augmentation in the mutant. This upregulation of BMP signaling in mutant cs was confirmed by immunostaining for pSmad1/5/9 (Fig 6I. J). In wt mice, BMP activation was restricted to the TTR, with minimal staining observed within the cs (Fig 6I). In contrast, the absence of the TTR resulted in close proximity of dura and cs, with a resulting increase in pSMAD1/5/9 staining within mutant cs (Fig 6J). In line with this observation, expression of BMP transcriptional targets Spp1 and Msx1 were found to be increased in mutant cs^29,30^ (Fig 6K, L, M, N). In summary, our data suggest a working model for the role of the TTR in mouse cs development. During normal development, the TTR secrets BMP antagonists and functions as a barrier to block pro-osteogenic signals derived from dura, to maintain patency of the coronal suture. The temporally regulated loss of the TTR, potentially through transdifferentiation, allows the precise regulation of the onset of cs differentiation in response to dura-derived cues. In the absence of the TTR, the dura is able to induce suture cell differentiation and ossification via increasing BMP signaling activity, resulting in premature suture closure (Fig 7).

**Figure 6.**
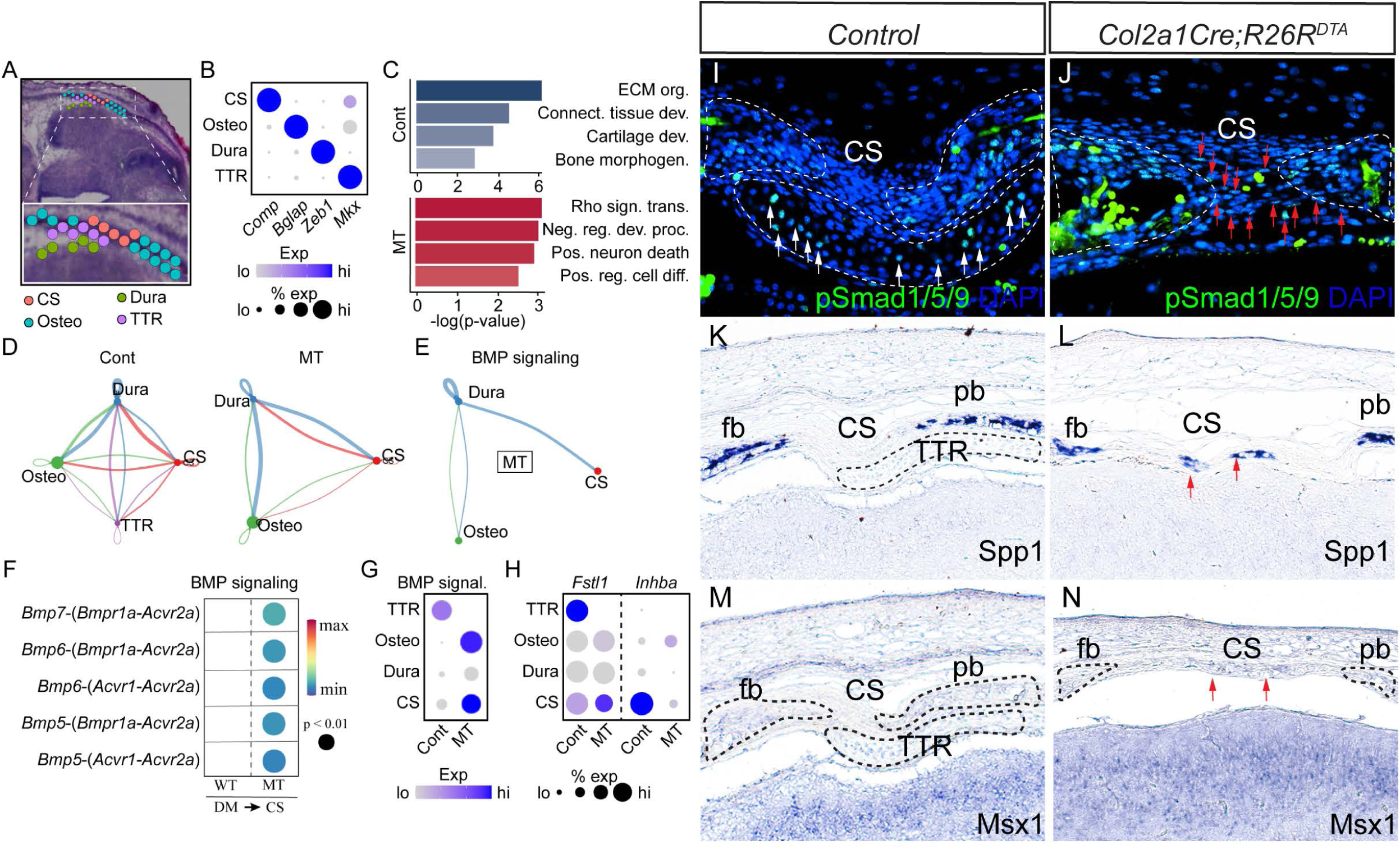
Spatial transcriptomics analysis of CS in littermate control and Col2a1Cre;R26R^DTA^ *mice*. (A) Sample overview shows regions of interest, including CS, Dura, Osteo and TTR. (B) Confirmed expression of spatially unique marker genes. (C) Pathway analysis of gene expression in CS of littermate control(*WT*) vs *Col2a1Cre;R26R^DTA^*(MT). (D) Overall communication strength *WT* vs *MT* (thicker line means more communication, line color denotes where signal is coming from). A stronger interaction between Dura and CS was observed in the *MT*. (E) BMP signaling is significantly strengthened in *MT*, while none was predicted in littermate control. (F) The ligand receptor pairs driving Bmp signaling in *WT* and *MT*. (G) Average expression of BMP signaling components and downstream targets. (H) Alterated expression of *Fstl1* and *Inhba* in *WT* and *MT*. (I, J) Immunostaining with anti-pSmad1/5/9 on transverse section of *WT* and *MT* at comparable level. Slides were counterstained with DAPI(blue). Blank arrows point to pSmad1/5/9 signal in WT TTR. Red arrows point to increased pSmad1/5/9 signal in CS of *MT*. (K-L) In situ hybridization with antisense probe of Spp1(K, L) and Msx1(M, N) on coronal sections of *WT* (K, M) and *MT* (L, N). Ectopic expression of Bmp signaling targets Spp1 and Msx1 in *MT* CS is marked by red arrows.

**Figure 7.**
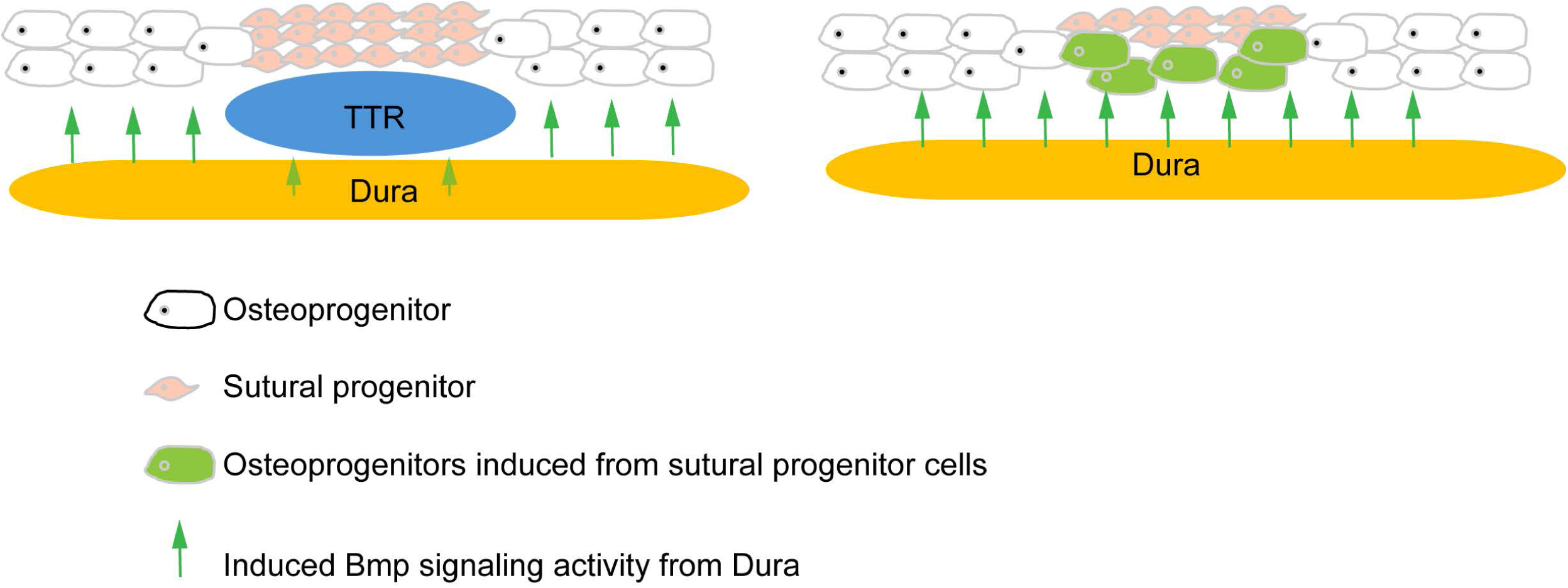
A hypothetical model illustrating the role of TTR in CS development.

## Discussion

The cranial endoskeleton is a cartilaginous structure of the mammalian skull formed prior to membranous ossification. Despite its close temporal and spatial association with the dermatocranium, its role in craniofacial development and malformation remains largely unknown. In this study, we focused on TTR, a temporary cartilage anlage located on the lateral wall of the cranial endoskeleton between coronal suture and the dura. We first examined the developmental process of TTR in a mouse model, characterizing its formation in relation to adjacent tissues, including calvarial bones and the coronal suture. Using a genetic approach to ablate TTR, we demonstrated that it is essential for maintaining coronal suture patency. Finally, through spatial transcriptomics, we further explored the mechanisms by which TTR regulates the coronal suture and surrounding tissues. Our findings suggest that TTR may modulate coronal suture development by antagonizing BMP signaling from the dura through tissue-tissue interactions. These results provide new insights into the role of the cranial endoskeleton.

### Function of transient cartilage in craniofacial development

Two types of cartilage tissues have been described in the chondrocranium^8^: some cartilages such as those in cranial base, undergo endochondral ossification; other cartilages, mostly located on the lateral wall of chondrocranium, exist transiently and are ultimately resorbed^31^. How the latter is implicated for normal craniofacial development remains largely unknown. TTR is a typical transient cartilage. Our data show TTR is formed around E13.5 and degrades by P5(Fig1, 2). It has been postulated that TTR determines the position of overlying suture^8^. In this study, we report that ablation of TTR causes premature fusion of coronal suture at E16.5. Our data suggest these transient cartilages play an important role in maintenance of coronal suture despite we cannot rule out the contribution by the shortening of cranial base to the coronal synostosis in current model. In consideration of the role of cartilage in tissue interaction, our findings are in line with previous report that the nasal cartilage is required for overlying transpalatal suture morphogenesis in rat^32^.

### Tissue-tissue interactions in suture morphogenesis

Tissue-tissue interaction plays a critical role in development. It orchestrates multiple processes such as cell differentiation, morphogenesis, and tissue patterning^33^. Temporal and positional juxtaposition of coronal suture and TTR indicate active interaction during development. Our data proved that TTR is required to maintain a patent coronal suture(Fig5). It will be interesting to test whether CS is required for normal TTR formation. A transgenic line driving Cre expression specifically in suture is ideal. In effort to identify a suture specific Cre tool, we have examined *Gli1CreER^T2^*, which has been used to study calvarial suture^34^. Unfortunately lacZ expression is detected in both CS and TTR(supplemental Fig2). Six2 is also a potential suture genetic marker^35^, and *Six2Cre* transgenic line labels certain cell population in embryonic craniofacial tissue^36^. However, we found Six2 is also expressed in the TTR (supplemental Fig3). Lineage tracing data show TTR is mostly derived from mesoderm cells, with a small percentage from neural crest(Fig3). Such a mesoderm/neural crest mixed pattern resembles to that of coronal suture, which develops closely adjacent to TTR^23^. Considering the same pattern and their close location, it would be interesting to explore the idea that TTR and CS share a common progenitor. Recent publications reported identification of suture specific genes using single cell RNAseq technique^35,37^. These are promising candidates to generate suture specific tools once their expression pattern is validated in embryonic suture tissues and exclusive from TTR. At same time, our spatial transcriptomics data can serve a good source to identify potential suture specific genetic markers(Fig 6).

### Interaction between dura, suture and cartilage

In *Col2a1Cre;R26R^DTA^* embryos, it is observed that the fusion of frontal bone and parietal bone always started from the basal level and close to the dura(Fig 5). This observation is in line with previous reports that dura plays an important role in suture formation^38-40^. Using in vitro rotation and translocation experiments, Bradely and colleagues showed that the “regional” posterior frontal dura influences cranial suture fusion in vitro^41^. This process is believed to be driven by inductive tissue interactions between dural cells and suture cells, mediated through growth factor signaling pathways. In case of coronal suture, TTR is located between coronal suture and dura(Fig 1). It thus might function as a physical barrier to prevent sutural cells inductive signals from the dura. Our data further show Bmp signal is significantly increased in CS and Osteo when TTR is removed, suggesting Bmp signaling is one candidate signal induced by dura. In addition to a function as a sole physical barrier, TTR also expresses Fstl, an antagonist of Bmp signaling (Fig 7H). In addition, our spatial data also show Fgf signaling is active in wildtype TTR and CS but is decreased in mutant CS (supplemental Fig4). Fgf is known to antagonize Bmp signaling in many developmental events^42-44^. In summary, our data suggest that TTR blocks Bmp signaling induction from dura by physical barrier and as a signal hub.

### Is TTR conserved between humans and mice?

Mice have been used as a model to study human diseases for decades. They are suitable tools to study birth defect because mouse development highly resembles human embryogenesis. Our present study showed TTR plays a crucial role in maintaining coronal suture during mouse development, and it raises a question whether there is a TTR counterpart in human fetus? Although it was previously suggested that humans lack TTR^45^, as most primates exhibit a significant gap between the ala orbitalis and parietal plate, a recent study utilizing chondrocranium scanning of a human fetus presented different findings^46^. In this report, researchers identified a pair of cartilage anlagen similar to the TTR observed in mice. Both these cartilages and TTR are located in the same region-the lateral wall of the developing chondrocranium between ala orbitalis and parietal plate-at comparable stages of gestation: week 13 in human fetuses and E13.5 in mouse embryos^47,48^. Additionally, these cartilages undergo morphological changes resemble mouse TTR, initially expanding before subsequently shrinking^46^. Using provided data, we have reconstructed an integrated model including both the dermatocranium and chondrocranium, and we observed that these cartilage anlages are located between frontal bone and parietal bone, same as TTR in mice (Fig.8). We thus cautiously suggest that the human fetus also has TTR and that in both mice and humans it may play a conserved role as we have demonstrated in the present study.

**Figure 8.**
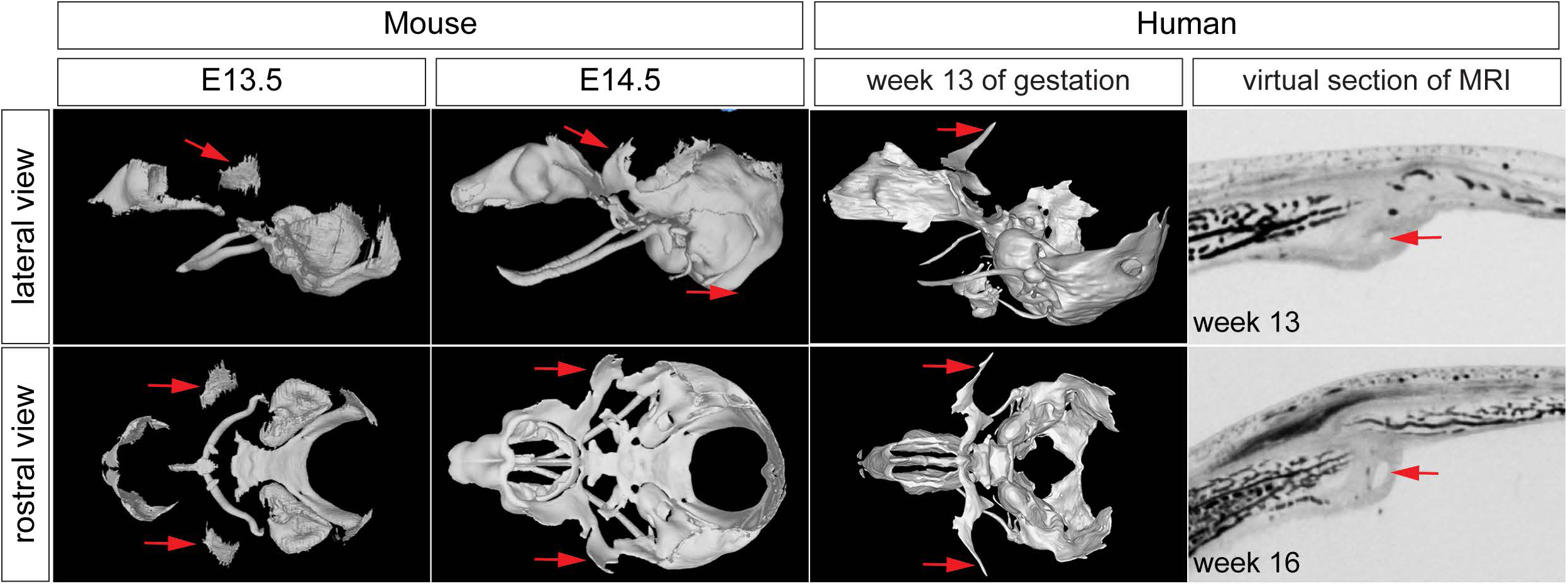
TTR is likely a conserved between mouse and human. (A, B, C, D) Scanning of mouse cranial endoskeleton at E13.5(A, B) and E14.5(C, D) is shown as lateral and rostral views^14,48^. (E, F) Scanning of human chondrocranium cartilage at week 13 of gestation at lateral(E) and rostral view(F). (G, H) Virtual section of MRI scanning of human fetus skull at week 13(G) and 16(H), respectively. Red arrow points to mouse TTR (A-D) and human counterpart(E-H). cs, coronal suture; fb, frontal bone; pb, parietal bone; TTR, tectum transversum.

## Materials and Method

### Mice

All animal experiments performed in the study were approved by the Institutional Animal Care and Use Committee of Tulane University. C57BL/6J mice were obtained from Jackson Laboratory (Bar Harbor, ME). The following transgenic mice used in this study were maintained on a C57BL/6J background: *Gt(ROSA)26Sor^tm1(DTA)Mrc^*designated here in as *R26R^DTA^* ^49^, and *Gt(ROSA)26Sor^tm1Sor^*referred to as *R26R^50^*. All other mice strain including *E2f1^Tg(Wnt1-cre)2Sor^*, designated as *Wnt1Cre2*^21^, *Mesp1^tm2(cre)Ysa^* abbreviated as *Mesp1Cre^22^*, and *Col2a1Cre^51^,* were maintained on a mixed *C57BL/6J; 129SvJaeSor* background. The vaginal plug was checked daily in the mating females and midday when vaginal plug was observed was considered as embryonic day (E) 0.5. For post-natal stages, the day on which pups were born was considered as post-natal day 0 (P0).

### Wholemount alcian blue staining

The whole mount alcian blue staining was performed according to previously published protocols^52^. Briefly, embryos of specific embryonic stages were dissected in PBS and then fixed in Bouin’s solution for overnight at room temperature. After several washes with 0.1% ammonium hydroxide in 70% ethanol for 24 hours, the embryos were equilibrated in 5% acetic acid for 2 hours. Further, embryos were stained with 0.05% alcian blue in 5% acetic acid for 2 hours. Following staining, embryos washed with 5% acetic acid for 2 hours and then cleared in methanol for 2 hours. Finally, embryo stored in benzyl alcohol/benzyl benzoate (BABB) solution.

### Skeletal preparation

The skeletal preparation was performed as published previously^53^. In short, E18.5 embryos or P5 pups were dissected and incubated in water for overnight at room temperature for 24 hours. Following removal of skin and evisceration, samples were fixed in 95% ethanol for overnight at room temperature. Further, samples were stained in alizarin red and alcian blue staining solution (0.005% alizarin red, 0.015% alcian blue and 5% glacial acetic acid in 70% ethanol) for 3 days at 37°C. After, briefly rinsing in 95% ethanol, samples were then cleared in 1% KOH solution for overnight and subsequently using the following series of glycerol/KOH solutions (20:80, 40:60, 60:40 and 80:20) each for 24 hours.

### Histology, alkaline phosphatase/alcian blue (AP-AB) and X-gal staining

Embryos were dissected in PBS and were fixed in 2% formaldehyde (from a 37% stock), and 0.2% glutaraldehyde (grade I, from a 25% stock) in PBS for 1 hour. Samples were then dehydrated in series of 15% sucrose and 30% sucrose for cryoprotection each for 24 hours, embedded in OCT blocks, and stored in −80°C. AP-AB staining was performed as described previously^54^. In short, slides were cryo-sectioned at a thickness of 10 µm, washed with PBS and incubated with 0.03% nitro-blue tetrazolium chloride (NBT) and 0.02% 5-bromo-4-chloro-3-indolyphosphate p-toluidine salt (BCIP) solution to observe AP activity in bone primordium. After rinsing with water, slides were then immersed in 1% alcian blue 8GX (A5268, Sigma) in 0.1 N HCl briefly to stain cartilage cells, and then counterstained with 0.1% nuclear fast red solution. For X gal staining, sections were dipped in standard staining solution (0.4% X-Gal, 5 mM potassium ferricyanide, 5 mM potassium ferrocyanide, 2 mM MgCl_2_, 0.01% sodium deoxycholate, and 0.02% Nonidet P-40 (NP-40) in PBS) for overnight at room temperature^18^.

### In situ hybridization and immunofluorescence staining

For *in situ* hybridization, E18.5 embryo were fixed in 4% paraformaldehyde (PFA) for overnight at 4 °C, proceeded to gradient ethanol dehydration and embedded in paraffin. The procedures for *in situ* hybridization was performed as described previously^53^. The immunofluorescence staining was performed on 10 μm frozen sections using standard protocol^55^. Briefly, sections were fixed with cold acetone for 10 min, washed in PBS with 0.1% Tween-20 (PBST) and blocked with 5% serum (goat or donkey) for 1 hour. The sections were then incubated in primary antibody diluted in PBS with 1.5 % serum overnight at 4°C. After washing in PBST, sections were exposed with Alexa fluor 594 or 488 conjugated secondary antibody (Thermo Fisher Scientific) in PBS with 1.5% serum (1:1000 dilution) for 1 hour at room temperature and mounted with VECTASHIELD antifade mounting media with DAPI (Vector laboratories). The following primary antibodies were used in the present study: anti-Runx2 (Cell Signaling Technology 12556, 1:200) anti-Sp7 (Abcam ab94744, 1:200), anti-ColX (Thermo Fisher 14-9771-82, 1:200), anti-cleaved Caspase 3 (Cell Signaling Technology 9664, 1:400) and pSmad1/5/9 (Cell Signaling Technology 13820, 1:200).

### 10X Visium spatial transcriptomics

Spatial transcriptomics analysis was carried out using the Visium Spatial Gene Expression System (10× Genomics, Pleasanton, CA) following the demonstrated protocol CG000239 for fresh frozen samples. In summary, E14.5 embryonic head samples were harvested, fresh-frozen in OCT, and stored at -80°C. Cryosectioning was performed at -25°C, and the sections were placed onto the capture areas of a Visium slide. Tissue fixation was achieved using methanol at -20°C, followed by dehydration with isopropanol and staining with hematoxylin and eosin (H&E). To determine optimal permeabilization conditions, enzymatic treatment was applied for varying durations (0–40 min), after which first-strand cDNA synthesis was performed using fluorescently labeled nucleotides. The slide was imaged using a Texas Red filter cube, and an optimal permeabilization time of 13 minutes was selected based on visual assessment to balance mRNA recovery and diffusion. For gene expression assay, library preparation, purification, and indexing followed standard protocols. Sequencing was conducted in a paired-end format using the Illumina NextSeq 2000 platform. Read alignment and demultiplexing were performed with the SpaceRanger pipeline using mouse genome reference (GRCm39). Seurat^56^ package was used for downstream analysis that included data quality control, normalization, integration and clustering. Differentially expressed genes were used as input for gene ontology analysis (GO) employing Enrichr tool web portal (https://maayanlab.cloud/Enrichr/)^57^. Cell-cell communication analysis between ligand-pair interaction used CellChat^58^. Module score analysis used Seurat’s *AddModuleScore* function and BMP genes identified from established KEGG pathways consistent with our previously publication^59^.

## Acknowledgements

We thank Dr. Mimi Sammarco for helping with 10X Visium spatial transcriptomics experiment. The authors are grateful for constructive comments from He lab members.

## Funding

This work was supported by NIH/National Institute of Children’s Health and Disease R01HD112474 to R.J.T. and by Tulane University and NIH/National Institute of Dental and Craniofacial Research grant DE028918 to F.H.

## Notes

### Competing Interest Statement

The authors have declared no competing interest.

