## Supplemental Figures for "The tectum transversum(TTR) maintains patency of the developing coronal suture"

**Supplemental Figure1. Apoptosis is not detectable in degenerating TTR.**

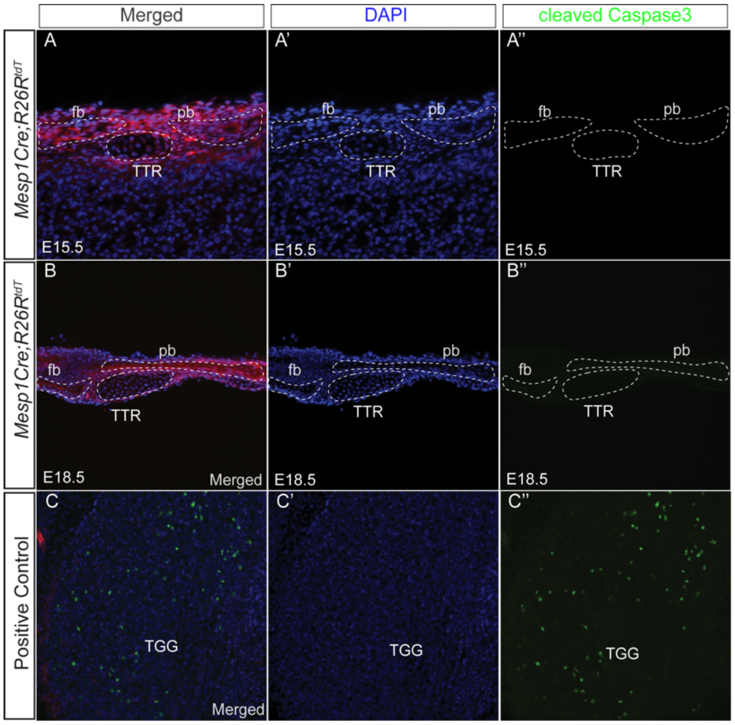


Supplemental Figure 1

Immunostaining of anti-cleaved caspase 3 on cross sections of *Mesp1Cre;R26R*$^{tdT}$ at E15.5(A, A', A'') and E18.5(B, B', B''). Trigeminal ganglia (TGG) is known to exhibit active apoptosis thus serves a positive control( C, C', C''). fb, frontal bone; pb, parietal bone; TTR, tectum transversum.

**Supplemental Figure 2: *Gli1CreER*$^{T2}$ labels cells in both CS and TTR.**

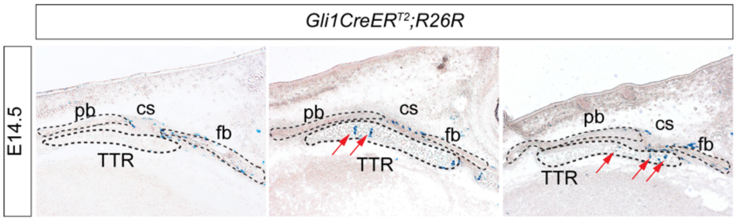


Supplemental Figure2

X-gal staining result of three *Gli1CreER$^{T2}$;R26R* embryos. LacZ expression (red arrows) is detected in TTR chondrocytes in two out of three embryos collected at E14.5, with tamoxifen administered at E8.5 by intraperitoneal injection to pregnant female. cs, coronal suture; fb, frontal bone; pb, parietal bone; TTR, tectum transversum.

**Supplemental Figure 3: Six2 is expressed in developing TTR and CS.**

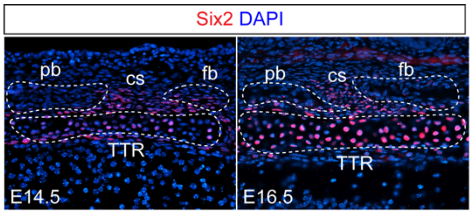


Supplemental Figure3

Immunostaining of Six2 in developing TTR at E14.5 and E16.5. Six2 expression is detected in both TTR and cs. cs, coronal suture; fb, frontal bone; pb, parietal bone; TTR, tectum transversum.

**Supplemental Figure 4: Alteration of major growth factor signaling in TTR ablation model.**

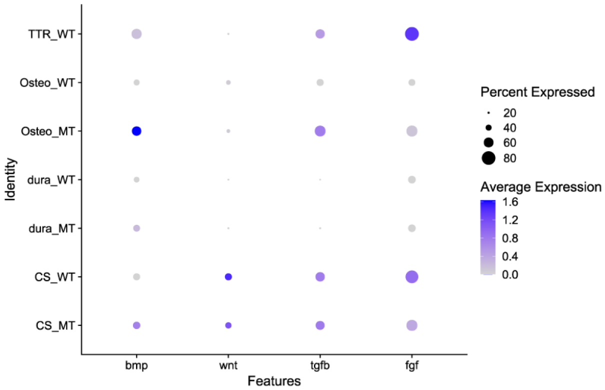


Supplemental Figure4

Examination of Bmp, Wnt, Tgfb and Fgf signaling activity in models with TTR and without TTR from 10X Visium spatial transcriptomics data.
